# Convolutions are competitive with transformers for protein sequence pretraining

**DOI:** 10.1101/2022.05.19.492714

**Authors:** Kevin K. Yang, Nicolo Fusi, Alex X. Lu

## Abstract

Pretrained protein sequence language models have been shown to improve the performance of many prediction tasks, and are now routinely integrated into bioinformatics tools. However, these models largely rely on the Transformer architecture, which scales quadratically with sequence length in both run-time and memory. Therefore, state-of-the-art models have limitations on sequence length. To address this limitation, we investigated if convolutional neural network (CNN) architectures, which scale linearly with sequence length, could be as effective as transformers in protein language models. With masked language model pretraining, CNNs are competitive to and occasionally superior to Transformers across downstream applications while maintaining strong performance on sequences longer than those allowed in the current state-of-the-art Transformer models. Our work suggests that computational efficiency can be improved without sacrificing performance simply by using a CNN architecture instead of a Transformer, and emphasizes the importance of disentangling pretraining task and model architecture.

## Introduction

Large pretrained protein language models have advanced the ability of machine learning models to predict protein structure and function from sequence. These models address the limitation that effective deep learning models generally require an abundance of labeled data to train. Since high-quality labels are only available for a limited number of sequences in most applications, protein language models first expose models to a large quantity of *unlabeled* sequences in a *pretraining* phase (Figure 1a), with the goal of imparting the model with a general foundation of knowledge about protein sequences so that they can be rapidly specialized to downstream tasks of interest with less training data than training from scratch (Figure 1b).

**Figure 1:**
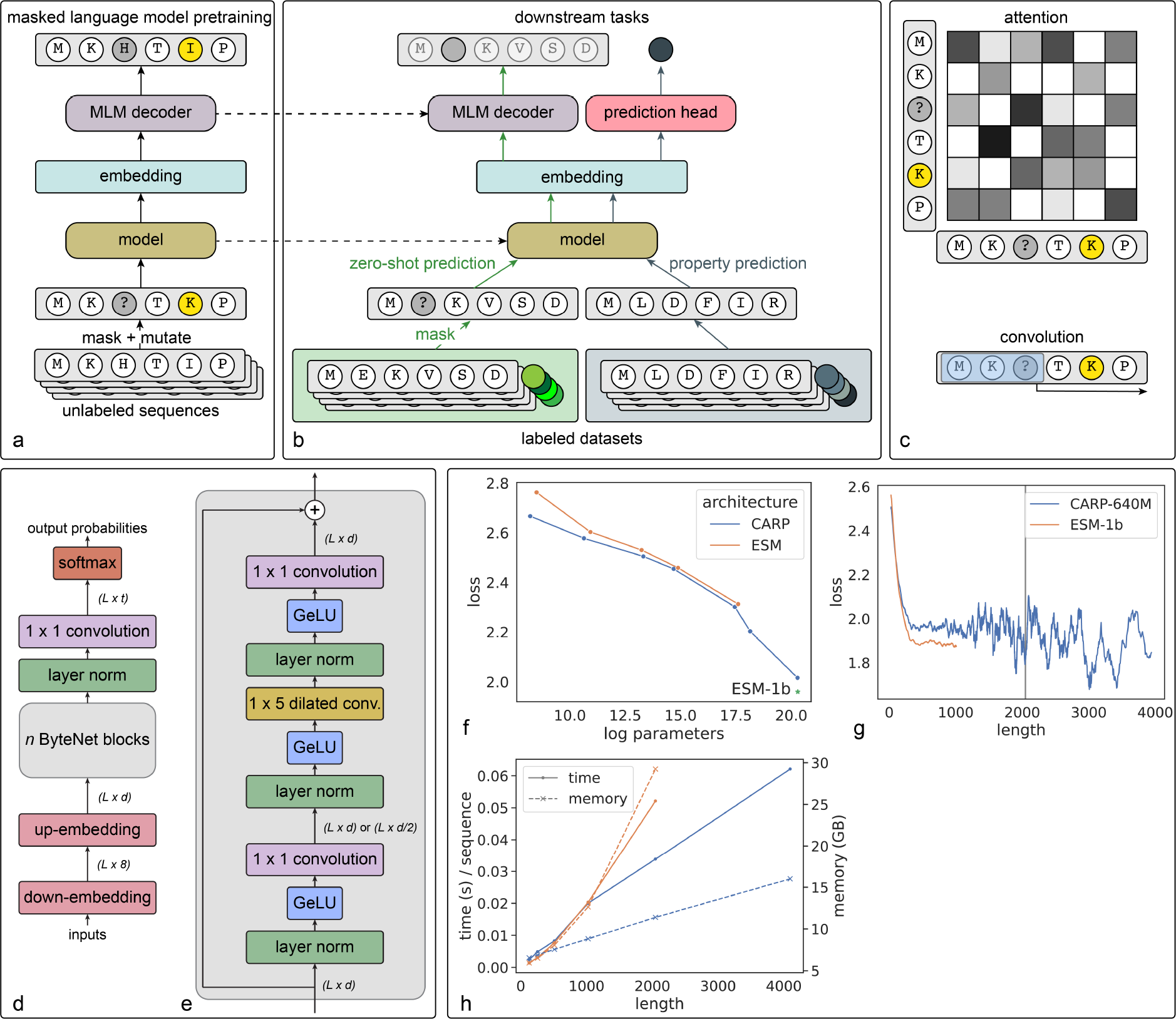
A schematic summary of our architecture, and pretraining, run-time, and memory performance of our model. a) Pretraining phase. Models are pretrained with masked language modeling: amino acids in unlabeled protein sequences are randomly masked and mutated to other amino acids, and the model is trained to recover the original sequence. b) After pretraining, models are specialized to downstream tasks. The weights of the pretrained model are transferred to the new task (dotted lines). For some zero-shot tasks like predicting the impact of mutations, the masked language modeling decoder is also useful and can be transferred. Otherwise, the decoder is replaced with a prediction head trained to output a prediction useful for the downstream task. c) Visual explanation of architecture options. Transformers parse sequences through self-attention modules (top), which are *L × L* matrices (requiring quadratic time and memory). CARP instead uses convolutions (bottom), which parse sequences with sliding window blocks, scaling linearly with sequence length *L*. d) CARP architecture e) ByteNet block f) Test loss on masked language pretraining task against number of parameters in model for CARP (blue) versus the state-of-the-art Transformer models (ESM - orange). The green asterisk shows the previously reported training loss of ESM-1b on their test dataset (which we did not retrain on our own dataset). g) Test loss on the masked language pretraining task for CARP-640 (blue) and ESM-1b against length of protein sequences. ESB-1b accepts a maximum of 1022-length sequences. CARP-640M is trained on a maximum of 2048 length sequences (vertical grey line), but can extend to arbitrarily long sequences during test time (we test up to length 4096). h) Runtime and memory for forward passes with different input sequence lengths for the CARP-640M (blue) and ESM-1b (orange) architectures, as measured using PyTorch [7] automatic mixed precision on a 48GB Nvidia A100 GPU.

While extensive effort has gone into validating protein language models for downstream applications, relatively little attention has been paid to the architecture of these models. Most works use a Transformer architecture, because these are the architectures employed in analogous work in natural language processing. However, Transformers have numerous drawbacks. First, the compute and memory requirements of these models scale quadratically with input sequence length. In principle, this is more of an issue in pretraining since back-propagating gradients requires additional memory, so longer sequences could be used during evaluation than during pretraining. However, a second limitation of Transformers is that because attention modules (Figure 1c) in these models are invariant to position, the position of each amino acid in a sequence must be encoded as part of the input. In most formulations, this position encoding is difficult to extend past the maximum length seen during training, and as a result, most popular pretrained language models limit the input length of sequences during both pretraining and inference. For example, ESM-1b and ESM-1v have a maximum length of 1022 residues. Unfortunately, this excludes many proteins of interest: of the 42 million cluster representatives in the March 2020 release of UniRef50 [1], 1.1 million, or 2.6%, are longer than 1022 residues, including the SARS-Cov-2 spike glycoprotein and the *Streptococcus pyogenes* CRISPR-associated endonuclease Cas9.

We reasoned that exploring alternative architectures could improve the computational efficiency of protein language models while allowing for a greater range of (longer) sequences to be studied. In this paper, we study if convolutional neural networks (CNNs) can be effective as pretrained protein language models. Unlike Transformers, CNNs scale linearly with input sequence size. Moreover, CNNs inherently incorporate relative positional information, since sequences are modeled as sliding windows of amino acids (rather than amino acids being treated as independent tokens in the Transformer framework) (Figure 1c). While no previous works have employed CNNs as large pretrained protein language models, prior studies have shown that CNNs are effective at predicting variant fitness for single protein families [2], annotating protein function [3], and in smaller-scale methods design studies [4], supporting our hypothesis that CNNs are effective for protein sequences.

We trained protein sequence CNN masked language models, which we refer to as CARP (**C**onvolutional **A**utoencoding **R**epresentations of **P**roteins). We show that CARP models are competitive with the current state-of-the-art Transformer model ESM [5, 6] on a variety of downstream prediction tasks, including structure prediction, zero-shot mutation effect prediction, and out-of-domain generalization on biologically-relevant protein engineering datasets. Because CARP scales linearly in computation with the input sequence and does not rely on an input positional embedding, it is straightforward to apply it to sequences longer than the longest sequences in training, which we demonstrate with zero-shot predictions of mutation effects in CRISPR-Cas9. Overall, the result that CNN models can perform as well as Transformers on downstream tasks has profound implications as these models are increasingly adopted for bioinformatics applications: our results suggest that tools built on large language models can improve their computational efficiency without sacrificing performance, simply by using a model with our CNN architecture over a Transformer. To faciliate these uses, we open-source and release weights and code for our CARP models.

## Results

### CARP achieves similar performance on the pretraining task to ESM

We trained a series of 7 CARP models with increasing numbers of parameters on nearly 42 million sequences from UniRef50. Each CARP model is analoguous to a Transformer protein masked language model, but with the Transformer layers replaced by ByteNet dilated CNN blocks [8], as shown in Figure 1d and e (see Supplementary Table S1 for details on parameters and architecture). Our largest CARP, CARP-640M, contains approximately 640M learnable parameters, comparable with the popular Transformer model ESM-1b’s 650 million parameters.

First, we sought to understand if our CARP architectures could solve masked language modeling as well as Transformer architectures. In this set-up, we are not yet assessing the effectiveness of these models on downstream function or structure prediction tasks, only how well the models can solve the pretraining task of predicting randomly masked or mutated amino acids in unseen sequences. There is no guarantee that performing well on the pretraining task necessarily translates to better performance on all downstream tasks of interest, as this requires that features that these models learn to extract from protein sequences to solve the pretraining task also be features useful to downstream tasks. However, at minimum, performing well on the pretraining task suggests that the models are learning effectively from the pretraining task. For example, if CNNs underperform Transformers on the pretraining task, this would suggest that CNNs are less effective at learning features from masked language modeling - we sought to rule out that this was the case.

We compared the loss of our CARP models against ESM Transformer models with similar numbers of parameters (Figure 1f) on our held-out test dataset. We note that ESM-1b trained and tested models on an earlier version of UniRef50 with with different train/test splits than our work. To produce a more fair comparison, we re-trained some ESM models using our dataset (orange line in Figure 1f), but because reproducing their largest model is too computationally expensive, we simply report results from ESM-1b on its test dataset (green asterisk in Figure 1f).

Overall, we observe that CARP’s performance on the pretraining task is comparable to ESM’s across several orders of magnitude of variation in the number of parameters when using the same pretraining dataset. Our largest model, CARP-640M, has a test loss of 2.02, comparable to ESM-1b’s loss of 1.96 on its test set.

Next, we asked how performance on the pretraining task interacts with sequence length. In principle, one advantage of CNNs is that they can scale to arbitrarily long sequences (including sequences longer than those seen during training), but for this advantage to be practical, the model must generalize to longer sequences. Figure 1g shows the masked language modeling loss by length for CARP-640M and ESM-1b on their respective test sets, smoothed with a window of 30 in the length dimension. For both models, the pretraining loss improves quickly until the sequence length reaches about 500 (shorter sequences are generally more challenging because they provide less context to reconstruct masked tokens), and then slowly thereafter. The maximum input length for ESM-1b during training is 1022, and this cannot be extended during test time. In contrast, for CARP-640M the maximum input length during training is 2048, but we calculate test losses for sequences with up to 4096 residues. Our results indicate that sequences with a length greater than 2048 have a comparable loss to sequences between 500-2048 length, suggesting that CARP-640M generalizes the pretraining task to sequences longer than those seen during training.

### CARP’s run-time and memory scale linearly with sequence length

Next, to confirm that our CARP models are more computationally efficient, we compared the run-time and memory requirements of the CARP-640M architecture against ESM-1b’s architecture at different sequence lengths (Figure 1h). Because the original ESM-1b model caps sequence lengths at 1022 residues, we randomly initialize a new ESM-1b model with a longer positional embedding (4096 residues). Although this new model is not expected to encode useful information for transfer to downstream tasks because it has not been pretrained, it serves as a good estimate of computational performance since run-time and memory is generally determined by the number of parameters and architecture, not the specific values of the parameters.

In general, we observe that while for smaller sequences, CARP-640M and ESM-1b have similar run-time (up to 1024 residues) and memory (up to 512 residues) requirements, ESM-1b scales both quadratically while CARP-640M scales linearly. As a result, due to memory limitations in our GPU, we are only able to evaluate sequences around 2048 residues with ESM-1b before an OOM (out of memory) error, while sequences of 4096 residues (and longer) are still possible on our GPU set-up with CARP-640M.

### CARP achieves comparable performance on downstream tasks to ESM

A key goal of protein language models is to encode information that can be rapidly transferred to improve performance on downstream prediction tasks (Figure 1b). Depending on the task, different methods for adapting protein language models are appropriate. *Zero-shot* methods do not require access to labels for further training, simply interacting with the pretrained model as is, and are well-suited for tasks where labels are too scarce to train a new model. Alternatively, if labels for a training dataset are available, a small neural network (a “prediction head”) can be built on top of representations from the pretrained model and trained to predict these labels. When training the prediction head, the pretrained model can be *frozen*, meaning that its parameters are not allowed to change and only the prediction head is trained, or *fine-tuned*, meaning the pretrained model’s parameters are trained jointly with the prediction head’s. Often, the decision of whether to freeze or fine-tune the pretrained model depends on how much training data is available: because more parameters are trainable when fine-tuning, there is a greater risk of overfitting.

To assess if CARP is capable of improving performance on downstream tasks, we curated a wide range of benchmarks, including predicting structure, the impact of mutations on fitness, and functional properties such as fluorescence, stability, or melting temperature. When training with downstream labels, we evaluate both freezing (“pt-fr”) and fine-tuning (“pt-ft”) our pretrained model, and compare against ESM Transformer models. Finally, to confirm that downstream performance is improved by the masked language modeling task, we provide a variety of baselines that do not use pretraining, including randomly-initialized weights (“na-fr” and “na-ft” in our tables), and linear ridge regression and a small CNN built on one-hot amino acid encodings of the protein sequences.

Finally, in addition to comparing against ESM, we also compare against Ankh [9] and ESM-MSA [10] to better understand how CARP compares against diverse modeling decisions in protein MLM work. Unlike CARP and ESM, which only use an encoder, Ankh uses an encoder-decoder architecture. While CARP and ESM embed single sequences, ESM-MSA embeds a multiple sequence alignment. We report performance across most downstream tasks in Table S7. Overall, we observe that while no model outperforms any other model at all or even most tasks, CARP performs comparably or better than most models across diverse downstream tasks.

### Protein structure

One of the most striking successes of protein language models is their ability to encode structural information without access to structural labels during pretraining. We evaluate CARP-640M’s ability to encode structural information through 3 tasks:

1. **Remote contact prediction** asks a model to predict whether the C_*β*_ atoms of two residues separated by at least 24 residues in the primary structure are within 8 Angstroms of other in the three-dimensional structure. We train on the trRosetta [11] training set and evaluate the precision of the top *L* predictions on the CAMEO hard [12] and CASP13-FM [13] test sets. For contact prediction, we downsample CARP embeddings to 128 dimensions, perform an outer product to produce 2-dimensional embeddings, and then pass that to a 24-layer dilated residual CNN based on the trRosetta architecture. This is the same as the procedure used by ESM-1b.
2. **Remote homology detection** asks a model to detect structural similarity across distantly-related sequences. We evaluate accuracy on the fold-level holdout set from TAPE [14].
3. **3-class secondary structure prediction** asks a model to predict whether each residue in a protein is part of a helix, strand, or other. We use the training and validation sets from TAPE and evaluate accuracy on the CB513 test set. For this task, we train a neural network consisting of two CNN layers, an LSTM, and a linear head on top of the pretrained model, as described in Rives et al. [5].

Since the structural tasks are computationally expensive, we do not fine-tune pretrained models for these tasks (and only train on top of frozen models), and only provide metrics for ESM-1b previously available in literature.

As shown in Table S5, pretraining improves performance for structure prediction tasks, and CARP-640M is competitive with ESM-1b. These results show that pretrained convolutions learn structural information from single sequences, just as pretrained transformers do.

### Zero-shot mutation effect prediction

Pretrained protein language models can predict experimental measurements of protein function without further training on sequence-fitness measurements or sets of evolutionarily-related sequences [15, 6]. Following Meier et al. [6], we score CARP-640M on 41 deep mutational scanning datasets originally compiled by Riesselman et al. [16]. These datasets measure the effects of thousands of mutations or combinations of mutations to a parent sequence. Details are described in Section 0.1.

Figure 2a compares zero-shot performance for CARP-640M, ESM-1b, ESM-1v, position-specific scoring matrices (PSSM), and ProtBert-BFD. ESM-1v results are for an ensemble of five transformers pretrained on UniRef90. Averaged across the 41 datasets, CARP-640M has a Spearman correlation of 0.49, compared to 0.46 for ESM-1b, 0.51 for ESM-1v, 0.46 for PSSM, and 0.43 for ProtBERT-BFD. CARP-640M outperforms ESM-1b on 22 out of 41 datasets, ESM-1v on 18 out of 41 datasets, PSSM on 26 out of 41 datasets, and ProtBERT-BFD on 25 out of 41 datasets.

**Figure 2:**
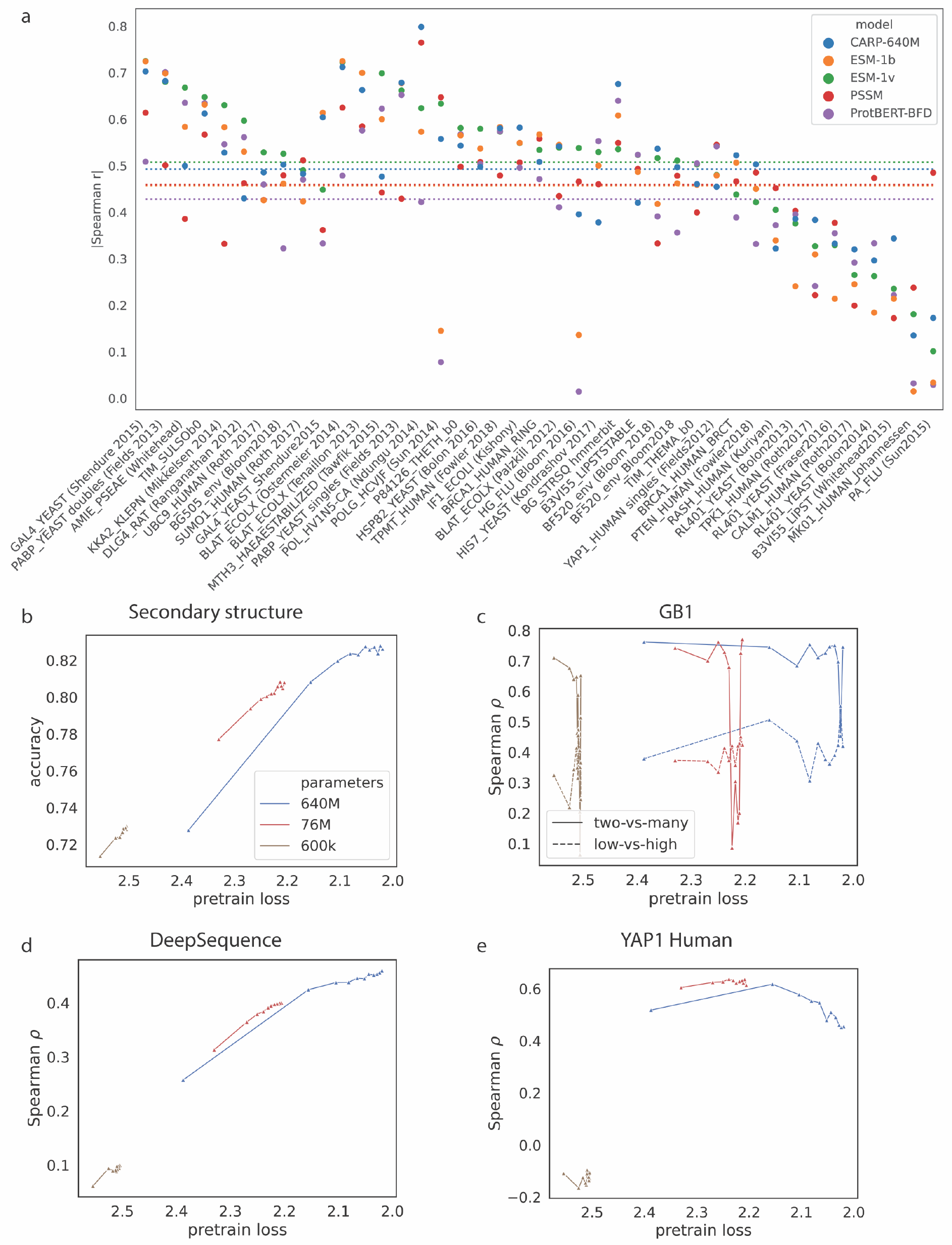
Performance on zero-shot fitness benchmarks and relationship between model size and loss on downstream performance on various tasks. a) Comparison of zero-shot fitness predictions across 41 deep mutational scanning datasets from DeepSequence. Points are Spearman correlation on each dataset. Horizontal lines show the average Spearman correlation across the datasets for each model. Values for ESM-1b, ESM-1v, PSSM, and ProtBERT-BFD are taken from [6]. b-c) Effect of model size and checkpoint pretrain loss on downstream performance for secondary structure (b) and GB1 (c). d-e) Effect of model size and checkpoint pretrain loss on zero-shot performance averaged over all DeepSequence datasets (d) and for YAP1 human (e)

Meier et al. [6] found that using the full UniProt sequences instead of only the sequence of the mutated domain results in better zero-shot predictions. However, this is not always possible with ESM-1x, as some UniProt sequences for these proteins are longer than 1022 residues. As a further proof of concept, we made zero-shot predictions for the effects of mutations in Cas9 from *Streptococcus pyogenes* [17], which is 1368 residues long, and obtain a Spearman correlation of 0.26. These results show that pretrained convolutions can make zero-shot predictions of protein mutation effects on fitness, including on sequences longer than allowed by ESM-1x.

### Out-of-domain fitness prediction

Another motivation for pretrained protein language models is that because they build on a general foundation of knowledge about proteins learned in the pretraining phase, they may be able to better extrapolate from limited training data. For example, a protein engineer may want to train a model on single mutants and make predictions for sequences with multiple mutations, or train a model that is accurate for sequences with fitness greater than what is seen in the training set. To test these kinds of out-of-domain prediction schemes, we evaluate CARP-640M on tasks from the following landscapes from FLIP [18].

1. AAV (Table S4): Adeno-associated virus (AAV) capsid proteins are responsible for helping the virus integrate a DNA payload into a target cell [19], and there is great interest in engineering versions of these proteins for gene therapy [20, 21]. [22] measure a rich mutational screening landscape for the VP-1 AAV protein, including 284,009 variants.
2. GB1 (Table S6): GB1 is the binding domain of protein G, an immunoglobulin binding protein found in Streptococcal bacteria. In their original study, Wu et al. [23] measured the fitness of 149,361 out of 160,000 possible combinations of mutations at 4 positions.

For each landscape, we evaluate several data splits where test sequences differ from training sequences:

- x-vs-many: Train on sequences with up to x mutations and test on the remainder of the landscape.
- mut-des: Train on sequences sampled from mutagenesis libraries and test on sequences designed by machine-learning models (AAV only).
- low-vs-high: Train on sequences with fitnesses below the wild-type and test on sequences with fitnesses above the wild-type.

In general, pretraining improves CARP-640M’s performance on these tasks, and fine-tuning the entire model outperforms freezing the pretrained weights. Comparisons to the baselines show that pretraining is most helpful when generalizing from single mutants to multiple. When not fine-tuning all the way through, there is little benefit from pretraining, and on some tasks pretraining hurts performance. CARP-640M outperforms ESM-1b on generalizing from few mutations to more, but ESM-1b is better at generalizing from a low-fitness training to higher-fitness sequences. These results show that pretrained convolutions help generalization to types of sequence variation not seen during training. On GB1, finetuning ESM-1b end-to-end instead of freezing the pretrained weights hurts its performance, while CARP-640M benefits from full finetuning. In addition, CARP-640M provides better representations without pretraining than ESM-1b on all the AAV tasks and 2 of the 4 GB1 tasks, showing that the architecture alone also influences generalization.

### In-domain property prediction

Finally, we consider property and fitness prediction tasks that do not require difficult biological generalization (Table S3). We evaluate on three sequence-fitness regression tasks:

1. **Fluorescence** requires the model to predict the effect of one or more mutations on the brightness of green fluorescent protein. The data was originally collected by Sarkisyan et al. [24]. We use the data splits provided in TAPE.
2. **Stability** requires the model to predict a small protein’s resistance to protease degradation. The data was originally collected by Rocklin et al. [25]. We use the data splits provided in TAPE.
3. **Meltome-mixed** requires the model to predict the melting temperature of a variety of proteins from across the domains of life. The data was originally collected by Jarzab et al. [26]. We use the cluster representatives and data splits provided in FLIP.

In addition, we evaluate on two intrinsically disordered region (IDR) function classification tasks taken from Zarin et al. [27]. For the IDR datasets, we use MMseqs2 [28] to cluster sequences to 50% identity and then randomly assign clusters to training, validation, or testing.

1. **Cdc28 binding** requires the model to predict whether an IDR is a target of Cdc28.
2. **Mitochondria targeting** requires the model to predict whether an IDR targets its protein for transport into the mitochondria.

In general, while pretraining improves CARP-640M’s performance on these tasks, neither of the large pretrained models consistently out-perform the baseline models on these tasks. Almost all the models perform very well on the IDR tasks, indicating that performance is saturating on these tasks. Nevertheless, CARP-640M is generally comparable to ESM-1b, showing that once again pretrained convolutions are comparable to pretrained attention.

### Evaluating performance at checkpoints illuminates relationship between pretraining and down-stream tasks

While we showed that our CNN CARP models perform comparably to ESM models across protein prediction tasks, not all downstream tasks benefit from pretraining with the masked language model task, with simple regression baselines performing as well as both CARP and ESM models in some tasks. One possibility is that for some tasks, the features learned to solve the masked language modeling pretraining task may not overlap or contain any useful information for some downstream tasks. We reasoned that one way to test this hypothesis would be to evaluate performance on downstream tasks at various checkpoints of our CARP models: if pretraining does learn features useful for downstream tasks, we expect a relationship where better performance on the masked language modeling task (which improves as models are pretrained for longer) has at least some correlation with performance on the downstream tasks.

We finetuned pretraining checkpoints for CARP-600k, CARP-76M, and CARP-640M on secondary structure, remote homology, and the FLIP GB1, AAV, and meltome tasks. Figures 2b and S2a show that structure prediction improves smoothly as the model size increases and the model is pretrained longer. This confirms that, as for transformers, pretraining imparts structural information to CNNs. However, Figures 2c, S2b, and S2c shows that this relationship does not exist for the out-of-domain FLIP tasks. In many cases, a small amount of pretraining is sufficient to outperform the naive baseline, and further pretraining has an unpredictable and often negative effect on performance. Finally, for the FLIP meltome task, Figure S2d shows that performance generally improves as CARP is pretrained, but the pretraining effect saturates, and CARP-76M outperforms CARP-640M.

Figure 2d shows that the average zero-shot performance improves with both model size and pretraining performance. However, this is not the case for every individual dataset within DeepSequence. Figure 2e shows a case where zero-shot performance peaks and then declines as CARP is pretrained. The Spearman correlation between the pretrain loss and zero-shot Spearman correlation range from 1 (monotonic increase in zero-shot performance with more pretraining) and -0.9, as shown in Figure S1. The average over the DeepSequence datasets is 0.40 for CARP-640M, 0.48 for CARP-76M, and 0.23 for CARP-600k. Although CARP-640M has better overall zero-shot performance than CARP-76M, CARP-76M more consistently improves with more pretraining than CARP-640M. The heterogeneity in the relationship between pretraining performance and zero-shot performance suggests that many but not all zero-shot tasks in DeepSequence are strongly determined by structural stability.

## Discussion

We show that CNNs can be comparable to or superior to Transformers on the masked language modeling pretraining task, and that pretraining helps CNNs achieve similar levels of performance as Transformers when adapting these models to downstream protein property prediction tasks. Our results challenge the tightly-held association between masked language modeling and Transformers, and shows that the pretraining task itself, not the Transformer architecture, is the essential component that makes pretraining effective. We believe that this insight is of practical utility to researchers working on proteins especially as protein language models are becoming a workhorse of bioinformatics methods. Unlike Transformers, CNNs scale linearly with input sequence length, which becomes important when modeling long protein sequences. Analogous work in natural language processing has also shown that CNNs can require fewer FLOPs of compute than Transformers, even for short sequences [29]. In addition, while we use standard dilated convolutions, there are more efficient convolution variants designed for sequence modeling [30] that may further improve model speed. Together, our work suggests that CNNs can improve the computational efficiency of prediction methods built on protein language models with no impact on performance, and further advances in these architectures, which are relatively underexplored compared to Transformers, may heighten this improvement.

One limitation of our CNN architecture is that unlike Transformers, it does not use a cross- or self-attention module. In at least some tasks, these modules can contribute towards interpretability. For example, it is possible to extract structural contact maps from pretrained Transformer self-attention matrices [31], and self-attention matrices contain information about binding sites [32]. Convolutions lack an obvious equivalent. In addition, attention-based models naturally extend to predict protein-protein interaction sites, because they provide a ready framework for pairwise interactions across amino acids between sequences. Finally, while we highlighted issues with computational efficiency of Transformers, recent technical advances have sought to address these challenges. For example, the challenge of quadratic dependence on sequence length can be ameliorated with approximate attention methods [33, 34, 35, 36, 37, 38, 39, 40] (although we note the choice of approximation matters for performance and the best method is not always clear *a priori* [41]). On proteins, [40] and [42] show that Performer approximate attention can perform well for autoregressive and masked protein language models, respectively, while ProteinBERT combines a fast global attention mechanism with masked language and functional annotation prediction pretraining [43].

However, we show that without pretraining, CNNs and Transformers perform differently on downstream tasks. This observation suggests that our CNNs may provide complementary inductive biases to those found in Transformer models. Unfortunately, we also find that, while masked language model pretraining is effective in imparting models with structural knowledge, the relationship between model size, pretraining loss, and downstream performance is less stable for out-of-domain protein engineering tasks, indicating that masked language modeling may not be effective for at least some types of tasks and emphasizing a need for more effective pretraining tasks. While we evaluate the effects of masked language model pretraining, numerous other pretraining tasks have been proposed including autoregressive language model pretraining [44], pairwise masked language modeling [45], and combining structural information [46, 47, 48, 49, 50, 51, 52] or functional annotations [43, 53]. Together, our work demonstrates the importance of disentangling pretraining task and architecture. We hope that this work is the first step in investigating the independent and interacting effects of pretraining and architecture for protein sequence modeling.

## Acknowledgments

We thank Brian Hie, Roshan Rao, Joshua Meier, and Alexander Rives for assistance with the zero-shot mutation effect prediction datasets.

## Author contributions

K.K.Y. conceptualized the project, wrote the software, performed the computational experiments, and wrote the initial paper draft. All authors interpreted the results and wrote the final paper.

## Methods

### Resource availability

#### Lead Contact

Further information and requests for resources and reagents should be directed to and will be fulfilled by the lead contact, Kevin K. Yang.

#### Materials Availability

No new materials were generated in this study.

#### Data and Code Availability

- Pretrained model weights and our train/validation/test splits for the two IDR datasets and UniRef50 have been deposited at https://doi.org/10.5281/zenodo.6564798 and are publicly available as of the date of publication. Other evaluation datasets used in this study are publicly available as of the date of publication. Repository links are listed in the key resource table.
- All original code has been deposited at https://github.com/microsoft/protein-sequence-models and is publicly available as of the date of publication. DOIs are listed in the key resources table.
- Any additional information required to reanalyze the data reported in this paper is available from the lead contact upon request.

### 0.1. Method Details

#### Architecture

One limitation of CNNs is that they are locally connected, so neurons in the model may not see the entire sequence at once, preventing the learning of distal interactions. To overcome this limitations, ByteNet uses a dilated CNN architecture [8], which increases the CNN perceptive field exponentially with the number of layers, allowing the model to obtain global context for long input sequences.

CARP combines the ByteNet encoder with simple input embedding and output decoding layers, as shown in Figure 1d and e. CARP begins with an embedding layer, which maps an input sequence of *L* tokens *x ∈* 𝔻^*L*^ to an 8-dimensional intermediate embedding, followed by a linear mapping into the model dimension *d*: *e*_0_ *∈* ℝ ^*L×d*^. This passes through a stack of *n* ByteNet dilated CNN blocks with residual connections in between followed by a final layer norm to produce the encoder representation *e*_*n*_ *∈* ℝ ^*L×d*^, and finally a linear decoder maps this to the *L × t* logits, where *t* is the number of possible tokens. The 1 *×* 5 convolution layer in every ByteNet block is dilated and padded to preserve sequence length. The CNN dilation rate doubles every layer up to a maximum rate *r* (for our experiments *r* = 128). This scheme is repeated multiple times in the network, always starting from a dilation rate of 1.

Throughout this paper, CARP refers to any ByteNet masked language model, while CARP-X refers to the model with approximately X parameters.) For example, our largest model, CARP-640M, has 640M parameters, comparable to the ESM-1b Transformer, which has 650 million parameters. Hyperparameters for different-sized versions of CARP and ESM are found in Tables S1 and S2, respectively.

### Dataset and Masked Language Modeling

We train CARP on the cluster representatives from the March 2020 release of UniRef50, with approximately 83k sequences held out for validation and another 210k sequences held out for testing, leaving 41.5 million sequences for training. Models are pretrained using the masked language model objective described in Rives et al. [5]. Each sequence is corrupted by changing some tokens to a special mask token or another amino acid token, and the model is tasked with reconstructing the original sequence. Specifically, 15% of tokens from each sequence are randomly selected. For those 15% of tokens, 80% are replaced by the mask token, 10% are replaced by a randomly-chosen amino acid, and 10% remain unchanged. The model is trained to minimize the cross-entropy loss between its predictions for the selected tokens and the true tokens at those locations.

### Training Details

We varied the number of parameters in CARP from approximately 3000 to 640 million by setting the model dimension *d*, setting the encoder hidden dimension *h*_*e*_ to either *d* or 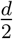, and setting the number of layers. All models are trained with the Adam optimizer, a maximum learning rate of 0.001, a linear warmup for 16,000 steps, and dynamic batching to maximize GPU usage. The largest model, CARP-640M, was trained on 128 32GB Nvidia V100 GPUs for 620,000 updates, or approximately 56 days.

To adapt CARP to downstream tasks, we use the output from the final layer norm in Figure 1 as the output representation. Unless otherwise noted, the prediction head consists of a learned attention that converts the output from *L × d* to *d* followed by a 2-layer neural network with hidden size *d*. For tasks with labels, we evaluate both freezing and fine-tuning CARP and compare to ESM-1b or ESM-1v. We finetune models with a maximum learning rate of 0.0001, a linear warmup over 1000 steps, and early stopping based on the validation set. Finetuning was performed on one 32 GB V100; depending on the task, finetuning took between several minutes to 48 hours. Where relevant, we also compare the CARP architecture with randomly-initialized weights, linear ridge regression, and the small CNN described in Dallago et al. [18] and Shanehsazzadeh et al. [54]. In our experiments fine-tuning models across checkpoints, we initialize the prediction heads with the same weights across all model checkpoints of the same size, to control for randomness between prediction heads.

### Zero-shot fitness prediction

For one-shot mutation impact prediction, we score sequences by masking every mutated position and computing the log odds ratio between the mutated and wild-type residues at each mutated position, assuming an additive model when a sequence contains multiple mutations:

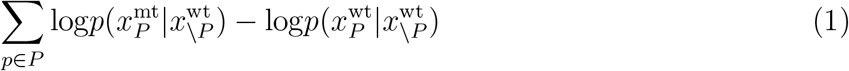

where *P* indicates the mutated positions.

## Supplemental Figure and Legends

### Hyperparameters for pretrained models of different sizes

All models are trained for 2 weeks on 1-8 32GB V100 GPUs with dynamic batching. Table S1 summarizes the hyperparameters for CARP. Table S2 summarizes the hyperparameters for ESM.

**Table S1:** CARP model hyperparameters. Max tokens is the maximum number of tokens per GPU per batch during training.

| Model | Parameters | Layers | $d$ | $d_{\text{MLP}}$ | Max tokens | GPUs |
| --- | --- | --- | --- | --- | --- | --- |
| CARP-640M | 643M | 56 | 1280 | 1280 | 11000 | $128 \times 32\text{GB V100}$ |
| CARP-76M | 75.7M | 32 | 1024 | 512 | 60000 | $16 \times 32 \text{ GB V100}$ |
| CARP-38M | 37.9M | 16 | 1024 | 512 | 40000 | $8 \times 32 \text{ GB V100}$ |
| CARP-24M | 23.9M | 16 | 256 | 128 | 400000 | $2 \times 32 \text{ GB V100}$ |
| CARP-600k | 608k | 16 | 128 | 64 | 600000 | $1 \times 32 \text{ GB V100}$ |
| CARP-40k | 415k | 16 | 32 | 16 | 600000 | $1 \times 32 \text{ GB V100}$ |
| CARP-4k | 3670 | 16 | 8 | 4 | 600000 | $1 \times 16 \text{ GB V100}$ |

**Table S2:**
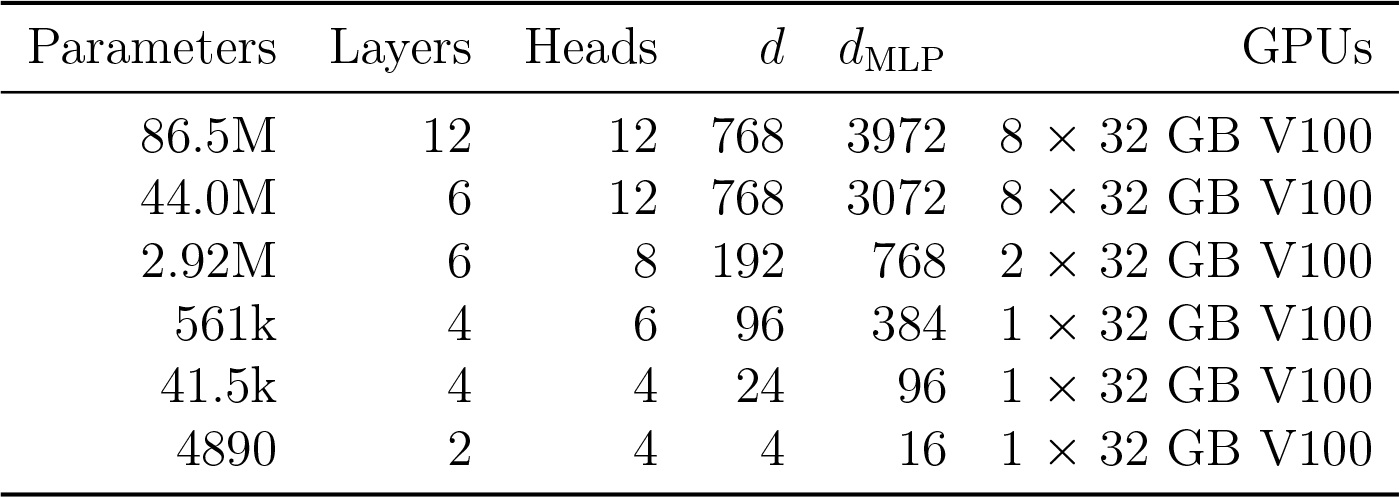
ESM model hyperparameters.

### Additional results

**Table S3:** Performance on in-domain tasks. For fluorescence, stability, and meltome, values reported are Spearman correlation. For the IDR tasks, values reported are area under the reciever operating curve. Values for ESM-1b on fluorescence and stability are taken from [5]. Values for baselines on fluorescence and stability are taken from FLIP. Uncertainties for ESM-1b and CNN are standard deviations over 10 random seeds except on the IDR tasks, where they are over 3 random seeds. Uncertainties for CARP-640M are standard deviations over 3 random seeds. We do not calculate uncertainties on meltome due to the computational cost.

| Method | Model | Task |  |  |  |  |
| --- | --- | --- | --- | --- | --- | --- |
|  |  | fluorescence | stability | meltome | Cdc28 | mito. |
| pt-fr | CARP-640M | 0.58±0.02 | 0.62±0.03 | 0.54 | 0.84±0.01 | 0.86±0.02 |
|  | ESM-1b | - | - | **0.67±0.01** | **0.91±0.004** | **0.90±0.01** |
| pt-ft | CARP-640M | **0.68±0.002** | **0.72±0.01** | 0.53 | 0.88±0.02 | 0.89±0.004 |
|  | ESM-1b | **0.68** | 0.71 | - | 0.89±0.01 | 0.88±0.01 |
| na-fr | CARP-640M | 0.62±0.01 | 0.52±0.17 | 0.29 | 0.84±0.01 | 0.86±0.01 |
|  | ESM-1b | - | - | 0.45±0.03 | 0.88±0.01 | 0.84±0.03 |
| na-ft | CARP-640M | 0.58±0.07 | 0.65±0.05 | 0.30 | 0.79±0.03 | 0.87±0.01 |
|  | ESM-1b | - | - | - | 0.83±0.02 | 0.85±0.02 |
|  | ridge | **0.68** | 0.48 | 0.17 | 0.52 | 0.53 |
|  | CNN | 0.67 | 0.51 | 0.34±0.01 | 0.84±0.02 | 0.87±0.02 |

**Figure S1:**
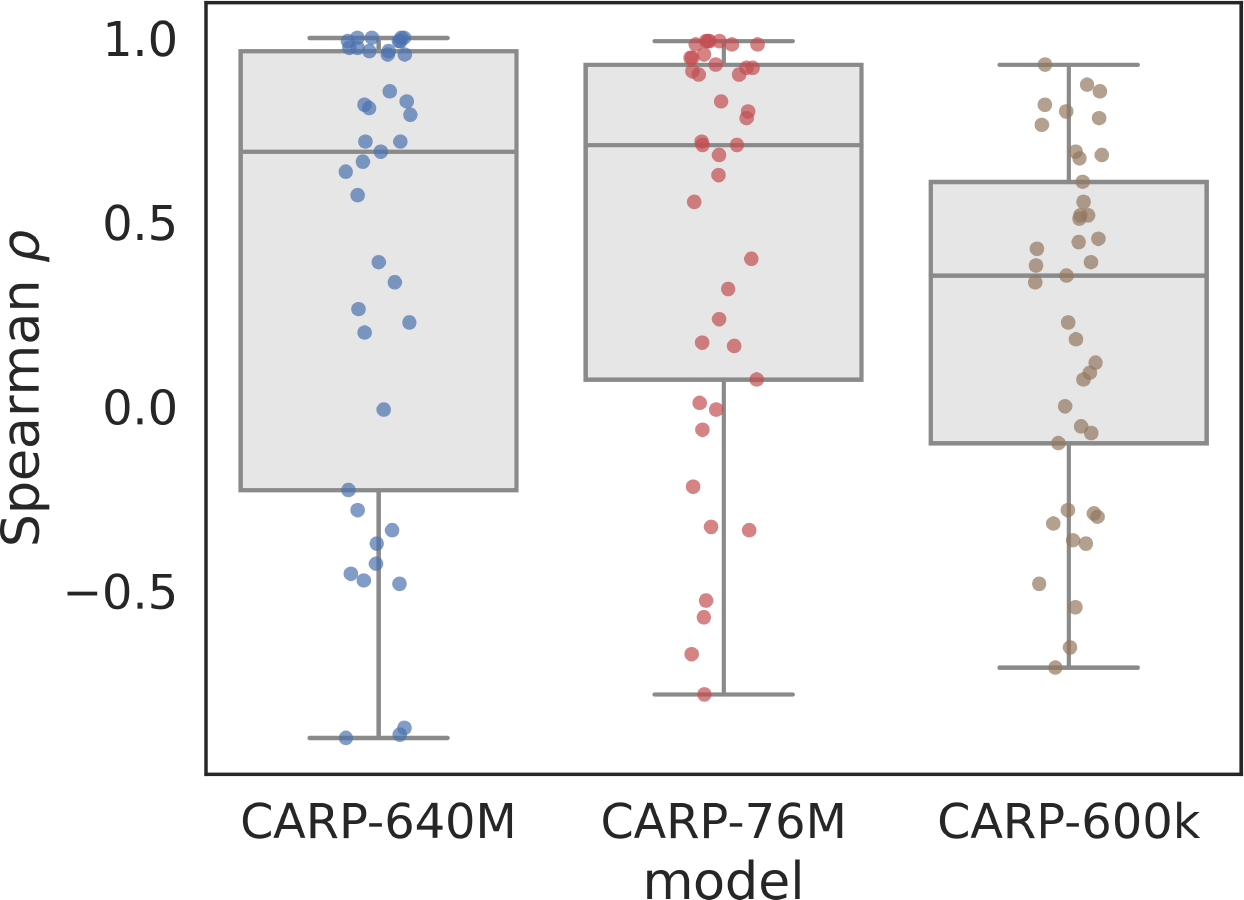
Spearman correlation between pretrained model checkpoint loss and zero-shot Spearman correlation across 41 deep mutational scanning datasets from DeepSequence.

**Table S4:** Performance (Spearman correlation) on the FLIP AAV tasks. Values for models other than CARP-640M are taken from [18]. Uncertainties for ESM-1b and CNN are standard deviations over 10 random seeds. Uncertainties for CARP-640M are standard deviations over 3 random seeds. [18] do not provide uncertainties for the mut-des task because of the computational cost.

| Method | Model | Task |  |  |  |  |
| --- | --- | --- | --- | --- | --- | --- |
|  |  | 1-vs-many | 2-vs-many | 7-vs-many | mut-des | low-vs-high |
| pt-fr | CARP-640M | 0.31±0.18 | 0.51±0.18 | 0.58±0.14 | 0.75±0.08 | 0.25±0.09 |
|  | ESM-1b | 0.03±0.11 | 0.61±0.04 | 0.65±0.01 | 0.76 | **0.38**±0.01 |
| pt-ft | CARP-640M | **0.73**±0.05 | **0.81**±0.03 | **0.77**±0.03 | **0.85**±0.003 | 0.19±0.08 |
| na-fr | CARP-640M | 0.48±0.07 | 0.50±0.05 | 0.60±0.05 | 0.76±0.02 | 0.21±0.02 |
|  | ESM-1b | 0.18±0.01 | 0.20±0.03 | 0.38±0.04 | 0.56 | 0.06±0.01 |
| na-fr | CARP-640M | 0.04±0.12 | 0.50±0.43 | 0.38±0.37 | 0.84±0.01 | 0.24±0.21 |
| baseline | ridge | 0.22 | 0.03 | 0.65 | 0.68 | 0.12 |
|  | CNN | 0.35±0.11 | 0.58±0.09 | 0.73±0.004 | 0.71 | 0.28±0.02 |

**Table S5:**
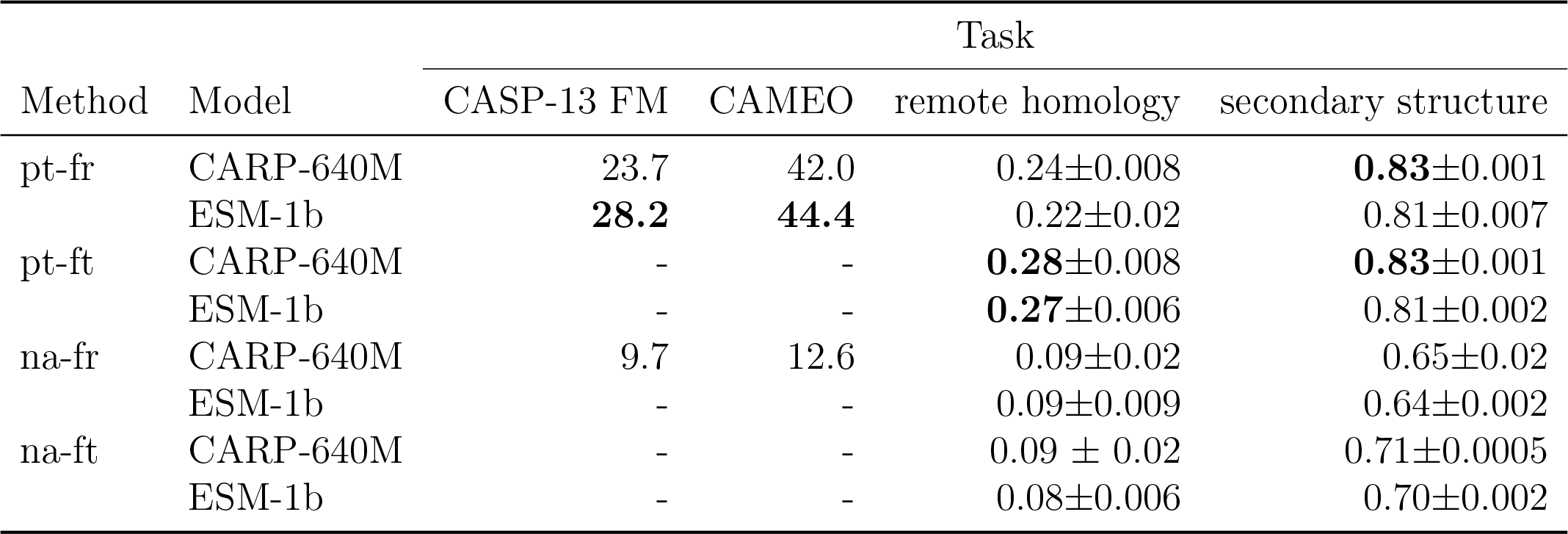
Structure prediction tasks. Values for ESM-1b on CASP-13 and CAMEO are taken from [10]. We report top *L* precision for CARP-13 FM and CAMEO and accuracy for remote homology and secondary structure. Uncertainties are standard deviations on 3 replicates with different weight initializations.

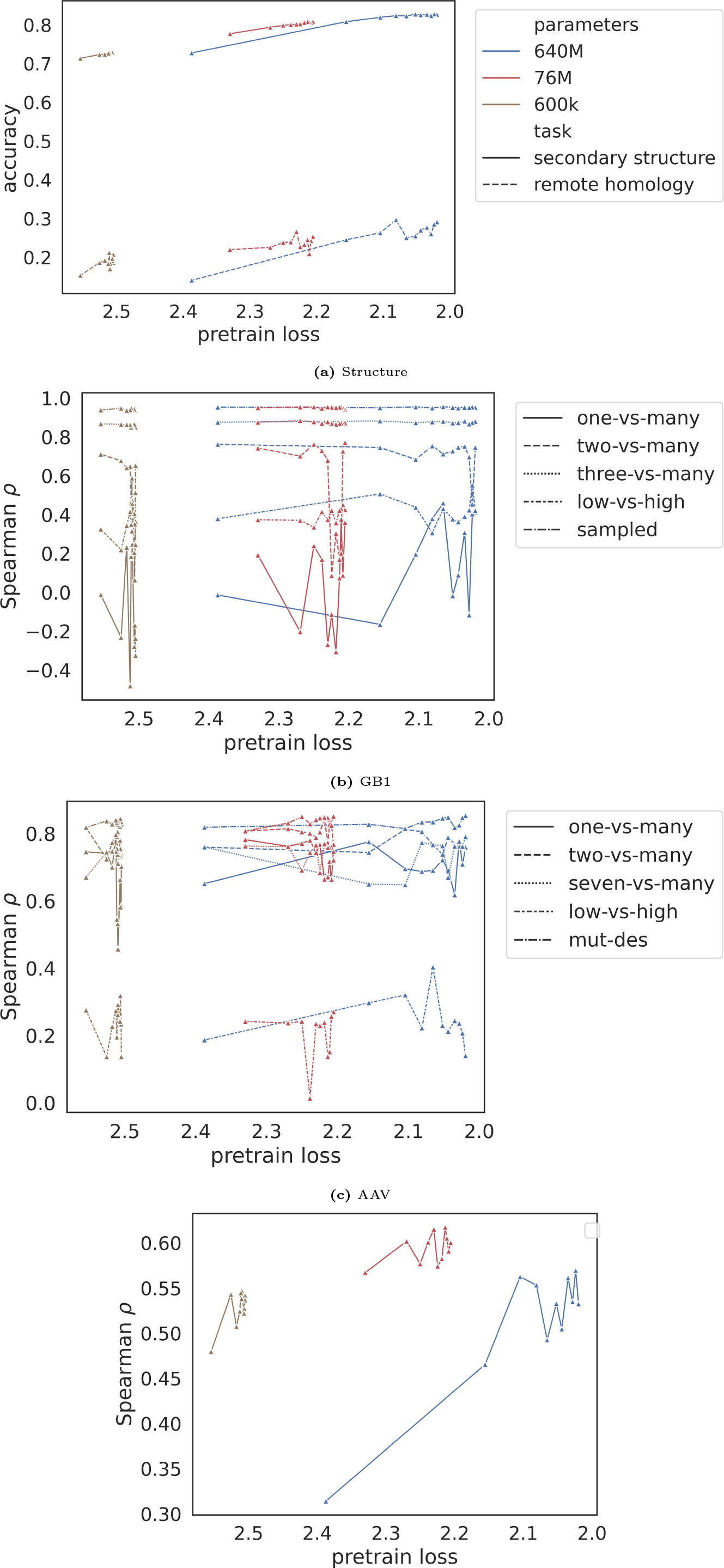

**Table S6:** Performance (Spearman correlation) on the FLIP GB1 tasks. Values for models other than CARP-640M and ESM-1b with full finetuning are taken from FLIP. Uncertainties for ESM-1b frozen and CNN are standard deviations over 10 random seeds. Uncertainties for CARP-640M and ESM-1b with full finetuning are standard deviations over 3 random seeds.

| Method | Model | Task |  |  |  |
| --- | --- | --- | --- | --- | --- |
|  |  | 1-vs-many | 2-vs-many | 3-vs-many | low-vs-high |
| pt-fr | CARP-640M | 0.15±0.18 | 0.18±0.23 | 0.62±0.06 | 0.12±0.03 |
|  | ESM-1b | **0.29**±0.02 | 0.47±0.05 | 0.79±0.01 | **0.53**±0.03 |
| pt-ft | CARP-640M | 0.19±0.26 | **0.73**±0.03 | **0.87**±0.004 | 0.43±0.04 |
|  | ESM-1b | 0.11±0.11 | 0.67±0.07 | 0.66±0.18 | 0.42±0.09 |
| na-fr | CARP-640M | 0.03±0.03 | 0.07±0.17 | 0.71±0.03 | 0.35±0.03 |
|  | ESM-1b | 0.12±0.01 | 0.21±0.01 | 0.52±0.01 | 0.32±0.03 |
| na-ft | CARP-640M | 0.11±0.07 | 0.38±0.26 | 0.68±0.33 | 0.23±0.26 |
|  | ESM-1b | 0.05±0.28 | 0.14±0.13 | 0.10±0.13 | -0.04±0.09 |
| baseline | ridge | 0.28 | 0.59 | 0.76 | 0.34 |
|  | CNN | 0.15±0.09 | 0.39±0.04 | 0.81±0.004 | 0.47±0.01 |

**Table S7:**
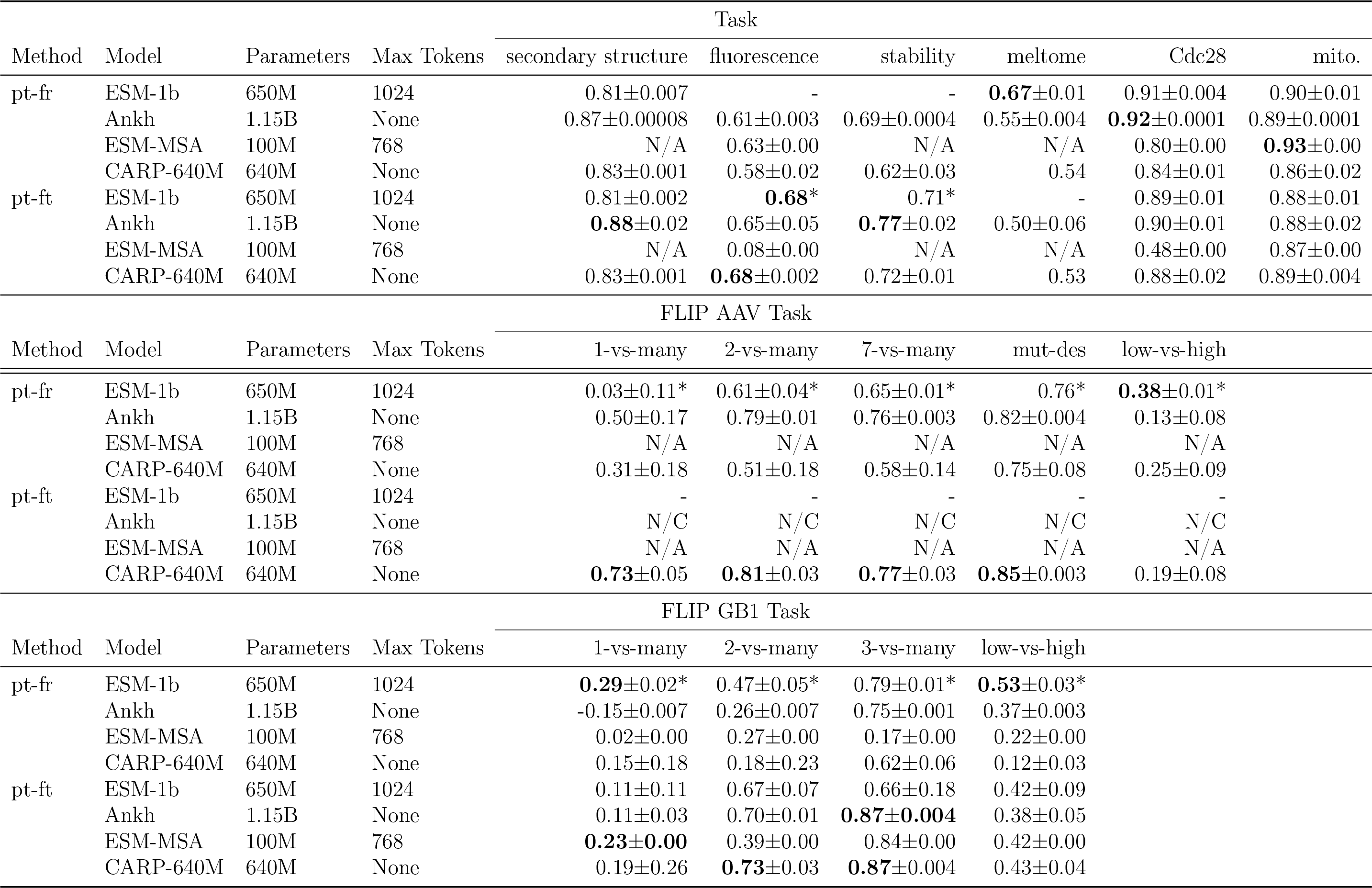
Performance on downstream tasks for ESM, Ankh, ESM-MSA and CARP-640M. For secondary structure we report accuracy. For the IDR tasks, values reported are area under the receiver operating curve. All other values are Spearman correlation. Uncertainties are standard deviations over 3 random seeds; if an experiment is missing uncertainty, replicates were skipped due to it being too computationally expensive. ^*^ denotes a value taken from [18]. - denotes an experiment that was skipped due to it being too computationally expensive. N/A denotes the model was not able to handle the benchmark (e.g. no MSAs for ESM-MSA). N/C denotes the model did not converge within 2 weeks of training.

